# The clathrin adaptor AP-1B independently controls proliferation and differentiation in the mammalian intestine

**DOI:** 10.1101/2023.05.12.540539

**Authors:** Maela Duclos, Anne Bourdais, Ophélie Nicolle, Grégoire Michaux, Aurélien Bidaud-Meynard

## Abstract

Maintenance of the polarity of the epithelial cells facing the lumen of the small intestine is crucial to ensure the vectorial absorption of nutrients as well as the integrity of the apical brush border and the intestinal barrier. Polarized vesicular trafficking plays a key role in this process, and defective transport due to mutations in apical trafficking-related genes has been shown to affect nutrient absorption. Interestingly, it has been demonstrated that downregulation of the polarized sorting clathrin adaptor AP-1B led to both epithelial polarity and proliferation defects in the mouse intestine. This enlightened a new function of polarized trafficking in the gut epithelium and a novel link between trafficking, polarity, and proliferation. Here, using CRISPR-Cas9-mediated mutation of the AP-1B coding gene *Ap1m2* in mouse intestinal organoids, we uncovered a novel proliferation pathway controlled by AP-1B. We showed that the polarity defects induced by *Ap1m2* mutations led to a defective apical targeting of both Rab11^+^ apical recycling endosomes and of the polarity determinant Cdc42. Moreover, we showed that these polarity defects were accompanied by an induction of YAP and EGFR/mTOR-dependent proliferation pathways. Finally, we showed that AP-1B additionally controlled a proliferation-independent differentiation pathway towards the secretory lineage. Overall, our results highlighted the pleiotropic roles played by AP-1B in the homeostasis of the gut epithelium.

## Introduction

Intestinal cells laying the gut lumen are constantly renewed by the treadmilling process of new cells arising from the proliferation of Lgr5^+^ intestinal stem cells (aka crypt-based columnar cells) and death of terminally differentiated cells at the top of the villus (Gehart and Clevers, 2019). The proliferation of intestinal stem cells (ISC) is controlled by niche factors emanating from both the neighbouring Paneth cells and mesenchymal cells secreting proliferative factors such as Wnt and the Wnt pathway agonist R-Spondin, as well as Epidermal Growth Factor (EGF) (Crosnier et al., 2006). Notably, varying exposure of cells in the transit-amplifying zone migrating towards the villus to niche gradients (e.g. Wnt, Notch) allows their differentiation into the different types of absorptive and secretory lineages of the small intestine (Beumer and Clevers, 2021). Among these cells, the major type is the enterocytes that bear a so-called brush border made of an array of microvilli increasing the membrane surface for nutrient absorption (Delacour et al., 2016).

The maintenance of the strong polarity of enterocytes is essential to ensure the vectorial transport of nutrients from the lumen to the blood vessels located in the underlying connective tissue. Polarized trafficking plays a major role in this maintenance and disturb trafficking to the apical membrane has been linked to major absorption defects, such as Microvillus inclusion disease (Schneeberger et al., 2018). Besides direct brush border defects, missense mutations in the gene coding a subunit of the AP-1 complex has been linked to an enteropathy through intestinal barrier defects (Klee et al., 2020). AP-1 is a clathrin adaptor tetrameric complex made of four subunits (two large subunits (β, γ), one medium subunit (μ1) and one small subunit (σ1)) (Nakatsu et al., 2014). Among them, the function of the epithelia-specific μ1B/μ2 subunit has been extensively studied in the basolateral targeting of membrane proteins (Fölsch et al., 2001; Gan et al., 2002; Gravotta et al., 2007). Furthermore, we and the group of Verena Gobel showed that AP-1 also mediates the sorting of apical cargoes in *C. elegans* intestinal cells (Shafaq-Zadah et al., 2012; Zhang et al., 2012), a function that has been confirmed in both cell culture and mouse models (Caceres et al., 2019; Hase et al., 2013). Notably, the polarized sorting defects induced by AP-1 subunits silencing resulted in the contralateral mislocalization of both transmembrane proteins and apical polarity determinants (e.g. Cdc42, PAR-6, PAR-3, aPKC) as well as in the apicalization of the enterocyte lateral membrane (i.e. ectopic lumen) (Hase et al., 2013; Shafaq-Zadah et al., 2012). Interestingly, it has been shown that *Ap1m2* knockout in mouse also induced a crypt hyperplasia, which enlightened a putative crosstalk between apical trafficking, cell polarity and proliferation in the gut. Here, we used CRISPR-Cas9-mediated mutation of *Ap1m2* in mouse intestinal organoids to study the relationship between cell polarity, proliferation, and differentiation in the mammalian intestine.

## Results and discussion

### *Ap1m2* mutations induce strong polarity defects in mouse intestinal organoids

In order to assess the role of AP-1B in intestinal cells integrity, *Ap1m2* gene, encoding the μ1B/μ2 subunit, was knocked down by conditional CRISPR-Cas9-mediated mutation in organoids (Bidaud-Meynard et al., 2022; Jad et al., 2022). Sanger sequencing of the genomic DNA of CRISPR-edited organoids using Inference CRISPR Edits software (Conant et al., 2022) showed that ∼82% of *Ap1m2* organoids were mutated, with half of the mutations inducing a single deletion of a guanosine located 5’ of the PAM site (Fig. 1A and Supplementary Fig. 1A). AP-1B is a renown sorting factor for basolateral proteins bearing dileucin or tyrosine-based signals (Fölsch et al., 1999) and has recently been involved in apical targeting (Caceres et al., 2019; Hase et al., 2013; Shafaq-Zadah et al., 2012; Zhang et al., 2012), making it a major player of apicobasal polarity in epithelia. To verify this in fully polarized enterocytes, the organoids culture medium was supplemented with valproic acid and inhibitor of Wnt protein 2 (IWP2), as described before (Bidaud-Meynard et al., 2022; Mosa et al., 2018). As expected, *Ap1m2* mutations induced a mispolarization of the brush border component phospho-ezrin, which accumulated at both apical and lateral membranes (Fig. 1B-C). This partial polarity inversion was also obvious by analysing the organoids by transmission electron microscopy (TEM), which revealed lateral ectopic lumen as well as regions of the enterocyte monolayer with a brush border located both at the apical and basal poles (Fig. 1D-F). In *C. elegans*, we found that these polarity defects are the consequence of the cytoplasmic dispersion of Rab11^+^ endosomes and a mistargeting of the polarity determinant Cdc42, that has been shown to be targeted to the plasma membrane through apical trafficking (Osmani et al., 2010). Mislocalised Cdc42 consequently recruited the other members of the Cdc42/PAR apical polarity module (PAR-3, PAR-6 and PKC-3) at the lateral membrane to convert it into an apical pole (Shafaq-Zadah et al., 2012). As ectopic lumens were observed upon *Ap1m2* mutation in organoids (Fig. 1E), we then assessed whether the same apical trafficking pathway is disrupted in organoids. Rab11 and Cdc42 immunostaining as well as PKCζ Immunohistochemistry (IHC) labelling showed a decreased apical confinement of all these factors and Rab11 endosomes were often detected at the lateral membrane (Fig. 1H-L). This suggested that the mechanisms by which AP-1B controls the apical polarity of intestinal cells are conserved in mammals. Moreover, TEM also revealed dilated Trans Golgi Network and abundant associated vesicles, multivesicular bodies and lysosomes (Supplementary Fig. 1B and Supplementary Table 1). Hence, these results confirmed the strong trafficking and polarity defects in *Ap1m2* CRISPR-edited organoids.

**Figure 1:**
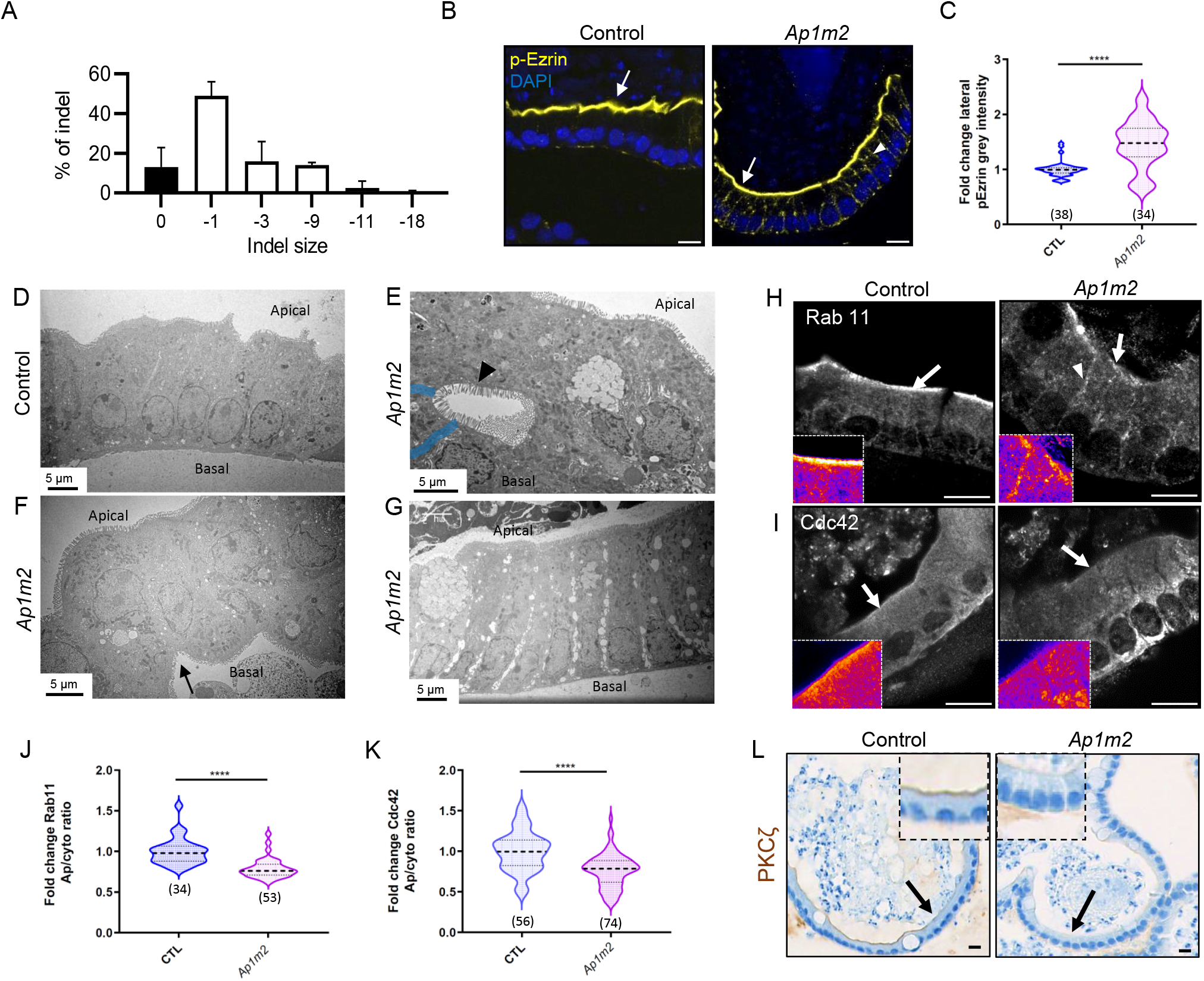
*Ap1m2* mutations induced polarity defects in fully differentiated organoids. (A) Analysis by Sanger sequencing of the genomic DNA of *Ap1m2* CRISPR-edited organoids (sg1 sgRNA). The histogram shows the mean ± SD of the repartition of indels found in the organoid population (N=2 independent experiments). (B-C) Phospho-Ezrin (p-Ezrin) staining in control or *Ap1m2* organoids. (B) shows representative images and (C) the quantification of the lateral p-Ezrin grey intensity, expressed in fold change (N=4 independent experiments). (D-G) Control and *Ap1m2* organoids were analysed by TEM. The arrow and the arrowhead show brush borders at the basal and lateral membranes, respectively. Lateral membranes in (E) are highlighted in blue. (H-K) Immunofluorescence staining of Rab11 endosomes (H) and Cdc42 (I) in Control (CTL) and *Ap1m2* organoids. Inserts in (H, I) represent fire-LUT coloured magnified images of the indicated apical ROIs (arrows). The arrowhead in (H) shows the lateral localization of Rab11. Panels (J-K) show the quantification of Rab11 and Cdc42 apical/cytoplasmic ratio in fold change (from 3 and 4 independent experiments, respectively). (L) IHC analysis of PKCζ in Control and *Ap1m2* organoids. The total number of organoids analysed is indicated in brackets, ^****^p<0.0001, Mann-Whitney test. Scale bars, 10μm (B, H-I, L) and 5μm (D-G).

### *Ap1m2* mutations induce an hyperproliferation in mouse intestinal organoids

In addition to apicobasal polarity, *Ap1m2* knockout in mouse additionally induced a crypt hyperplasia (Hase et al., 2013). To confirm this in our model, the proliferation of intestinal cells was measured using Ki67 staining in organoids cultured in normal EGF/Noggin/R-spondin1 (ENR) medium (i.e., presenting both crypt and villus-like regions) (Sato et al., 2011). *Ap1m2* mutations induced a 2-fold over-proliferation of intestinal cells compared to control organoids (Fig. 2A-B). This hyperplasia was correlated with a stacking of the cells which became thinner and longer (Figs 1G and 2C). The enterocyte monolayer was also dramatically disturbed, with a misalignment of nuclei and intertwined cells (Figs 1F and 2C). Furthermore, we observed intercellular junctions defects with many holes between cells (Figs 1G and 2C) as well as disturb adherens and tight junctions organization (Fig. 2D), as already reported upon AP-1 complex subunits downregulation (Hase et al., 2013; Klee et al., 2020). However, while cell-cell junctions were clearly affected, we were unable to observe the dotted cytoplasmic accumulation of E-Cadherin (Hase et al., 2013). This data favoured a pathway in which E-Cadherin intracellular displacement freed β-catenin that accumulated intracellularly and activated downstream proliferation targets (Hase et al., 2013). Furthermore, while Wnt pathway inhibition with IWP-2 completely effaced the crypts to only leave differentiated enterocytes and goblet cells in control organoids, crypt-like structures and Ki67^+^ proliferating cells persisted in *Ap1m2*-mutated organoids (Fig. 2F-H and Supplementary Fig. 1C-D). This suggests that the activation of the β-catenin pathway described earlier might still rely on mesenchymal Wnt signals, which are very important for small intestine proliferation and homeostasis in the context of the organ (Valenta et al., 2016), and that additional intestine-autonomous proliferation pathway(s) may be induced upon *Ap1m2* mutations. Moreover, we observed that the polarity defects induced by *Ap1m2* mutations, such as the formation of ectopic lateral lumen and the contralateral mislocalization of the apical and basolateral proteins P-ezrin and EphrinB2, respectively, were also observed in proliferating organoids (Fig. 2I-K). Considering that both *Cdc42* and *Rab11* knockout also induced an intestinal cells hyperplasia in mice (D’Agostino et al., 2019; Melendez et al., 2013; Zhang et al., 2022), these results raised the possibility that *Ap1m2* mutations induced an additional, mesenchyme-independent, proliferative pathway that could be linked to Cdc42 polarity defects.

**Figure 2:**
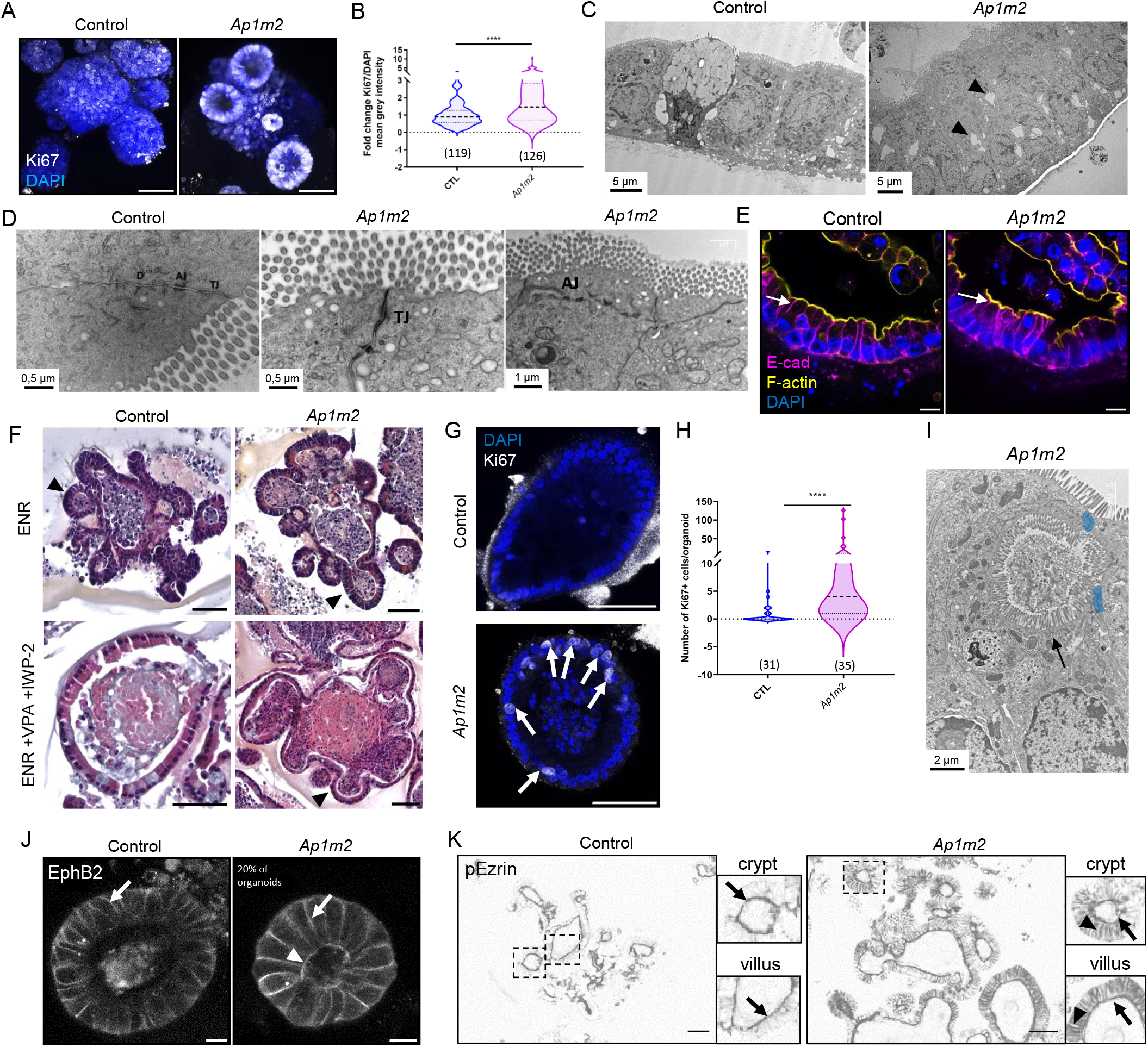
*Ap1m2* mutations induced an hyperproliferation in mouse SI organoids. (A-B) Immunofluorescence staining of Ki67 in Control (CTL) and *Ap1m2* organoids. (A) shows representative z-projections and (B) the quantification of Ki67/DAPI mean grey intensity ratio, expressed in fold change (N = 11 independent experiments). (C-D) TEM images of normally grown proliferative Control and *Ap1m2* organoids. Arrowheads highlight the holes between cells. (D) shows a focus on cell-cell junctions. D, desmosome; AJ, adherence junctions; TJ, tight junctions. (E) Immunofluorescence staining of E-Cadherin (magenta), F-actin (yellow) and Dapi in Control and *Ap1m2* organoids. Arrows indicate cell-cell junctions. (F) Haematoxylin-Eosin-Saffron (HES) staining of control and *Ap1m2* organoids cultured in normal ENR or full-differentiation (ENR+VPA+IWP-2) mediums. Arrowheads show intestinal crypts. (G-H) Immunofluorescence staining of Ki67 in fully differentiated Control (CTL) and *Ap1m2* organoids. (H) shows the quantification of Ki67^+^ cells/organoid. (I) *Ap1m2* organoids cultured in normal ENR also present ectopic lateral lumen (arrow), analysed by TEM. Lateral membranes are highlighted in blue. (J) Immunofluorescence staining of Ephrin B2 in control and *Ap1m2* organoids. Arrows and the arrowhead indicate the basolateral and the apical membranes, respectively. (K) IHC staining of p-Ezrin in control and *Ap1m2* organoids. Arrows show the apical membrane and arrowheads show the basolateral accumulation of P-ezrin. Inserts highlight the crypt and villus regions indicated by dashed lines. The total number of organoids examined is indicated in brackets, ^****^p<0.0001, Mann-Whitney test. Scale bar, 50μm (F, G, K) or 10μm (E, J).

### *Ap1m2* mutations-induced hyperproliferation involves YAP and EGFR/mTOR

To test this hypothesis, we stained Cdc42 in organoids edited with two different sgRNAs targeting *Ap1m2*. We found that Cdc42 apical localization also decreased in normally cultured, proliferating organoids (Fig. 3A). It has recently been shown that Rab11 is involved in the targeting of the Hippo signalling factor Yes-associated protein (YAP), and that *Cdc42* silencing in mouse intestine induced an hyperplasia through YAP-EGF-mTOR signalling (Goswami et al., 2021; Zhang et al., 2022). Notably, this pathway led to an initial increase of nuclear YAP expression by inhibiting its ubiquitin-dependent proteolysis (Zhang et al., 2022). Interestingly, YAP nuclear expression in crypts increased by 1.5-fold upon *Ap1m2* mutations, which indicated that the YAP-EGFR-mTOR pathway could be enhanced (Fig. 3B-C). To verify this, the proliferation of intestinal cells was measured upon treatment of the organoids with the EGFR and mTOR signalling inhibitors Afatinib and Rapamycin, respectively. Remarkably, both treatments rescued the proliferation of *Ap1m2*-mutated organoids to control levels (Fig. 3D-E) which confirmed that AP-1B also controls intestinal cells proliferation through YAP, EGFR and mTOR. Surprisingly, while *Cdc42* knockout has been proposed to induce EGFR and mTOR proliferative pathways as a consequence of an increased expression of YAP (Zhang et al., 2022), the accumulation of YAP was also cancelled by Afatinib and Rapamycin treatment (Fig. 3F-G). This suggests either the existence of positive feedback loops, or that YAP acts downstream of the mTOR pathway, as proposed before (Liang et al., 2014) or, finally, that YAP and EGFR pathways are induced in parallel. Indeed, in addition to YAP, Rab11 has been shown to mediate the membrane localization of Merlin, a YAP and cortical actin regulator that has also been involved in the control of EGFR signalling (Curto et al., 2007; Goswami et al., 2021). Furthermore, while Afatinib but not Rapamycin also slightly decreased the proliferation of control organoids by 1,5-fold, it dramatically inhibited *Ap1m2*-mediated overproliferation (∼8.3 fold) (Supplementary Fig. 1E-F). Hence, these results suggest that *Ap1m2* mutations lead to an hyperproliferation of intestinal cells by displacing Cdc42 from the apical membrane and activating YAP and EGF-mTOR-dependent proliferation pathway(s).

**Figure 3.**
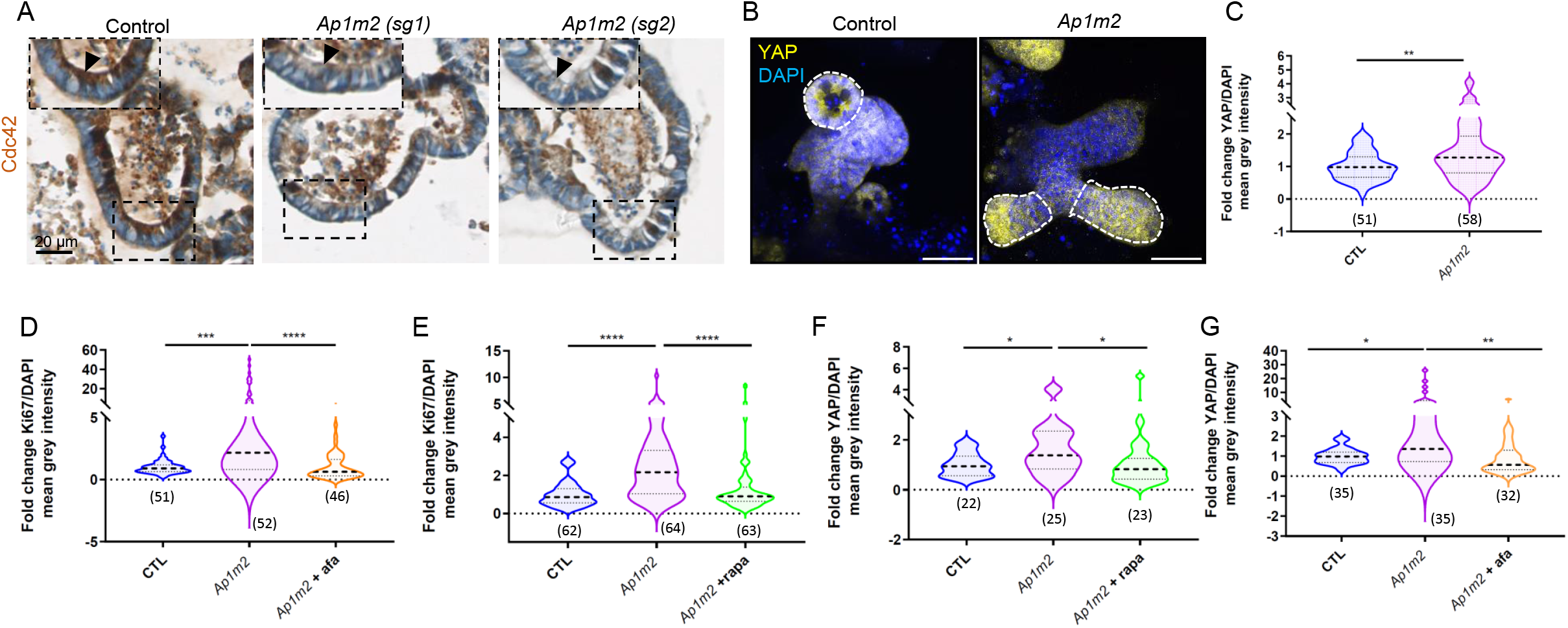
*Ap1m2* mutations induced a Cdc42, YAP and EGFR/mTOR-dependent hyperproliferation. (A) IHC staining of Cdc42 in control and *Ap1m2*-mutated organoids. sg1 and sg2 indicate organoids where *Ap1m2* has been edited with two different sgRNAs. The arrowheads show the apical membrane in the inserts, that are magnified images of the indicated ROIs. (B-C) Immunofluorescence staining of YAP in control (CTL) and *Ap1m2* organoids. (B) is a representative z-projection and (C) is the quantification of the nuclear YAP/DAPI mean grey intensity ratio expressed in fold change (N=7 independent experiments). (D-G) Quantification of Ki67/DAPI (D-E) and YAP/DAPI (F-G) ratios in control (CTL) and *Ap1m2* organoids treated with 200nM Afatinib (afa) (N=6 independent experiments) or 1 μM Rapamycin (rapa) for 3 days (N=7 independent experiments). The total number of organoids analysed is indicated in brackets, ^*^p<0,05, ^**^p<0,01; ^***^p<0,001; ^****^p<0,0001, unpaired Mann-Whitney test. Scale bar, 50μm.

### *Ap1m2* mutations induced major and proliferation-independent differentiation defects

Besides the defects observed in the arrangement of cells within the monolayer and in the enterocytes’ polarity, we observed that *Ap1m2* mutations induced severe differentiation defects. Indeed, Olfm4^+^ ISCs and Lysozyme^+^ Paneth cells, which are normally confined in the crypt, invaded the villus region of *Ap1m2*-mutated organoids (Fig. 4A-D). Furthermore, TEM analysis of the secretory cells revealed the presence of cells with a mixed zymogen/mucigen content - while the number of normal goblet and Paneth cells remained unchanged - upon *Ap1m2* gene editing (Fig. 4E-K and Supplementary Table 1), thus demonstrating a strong defect in secretory lineage differentiation. Interestingly, this rare phenotype, together with the invasion of the villus by Paneth cells, were also observed upon *Cdc42* knockout in mouse (Melendez et al., 2013). Of note, the invasion of the villus by Paneth and ISCs was not affected by Rapamycin nor Afatinib treatment, which indicated that these differentiation defects are independent of the YAP/EGF/mTOR proliferation pathways induced by *Ap1m2*-mutations (Fig. 4L-M). These results corroborate previous data showing that inhibition of EGFR signalling affected the proliferation but not the differentiation into goblet and Paneth cells (Basak et al., 2017). Further studies would be required to understand how AP-1B controls the differentiation of the secretory lineage in the intestine. This might rely on an imbalance of Wnt and Notch signals, which are crucial in determining the fate of secretory cells (Beumer and Clevers, 2021), and whose receptors’ membrane targeting is controlled by the AP-1 complex (Benhra et al., 2011; Wieffer et al., 2013). Overall, our results highlight the major roles played by AP-1B in the homeostasis of the small intestine cells, by regulating polarized trafficking, proliferation as well as differentiation (Fig. 4N) and confirm that AP-1B is essential for apical trafficking.

**Figure 4:**
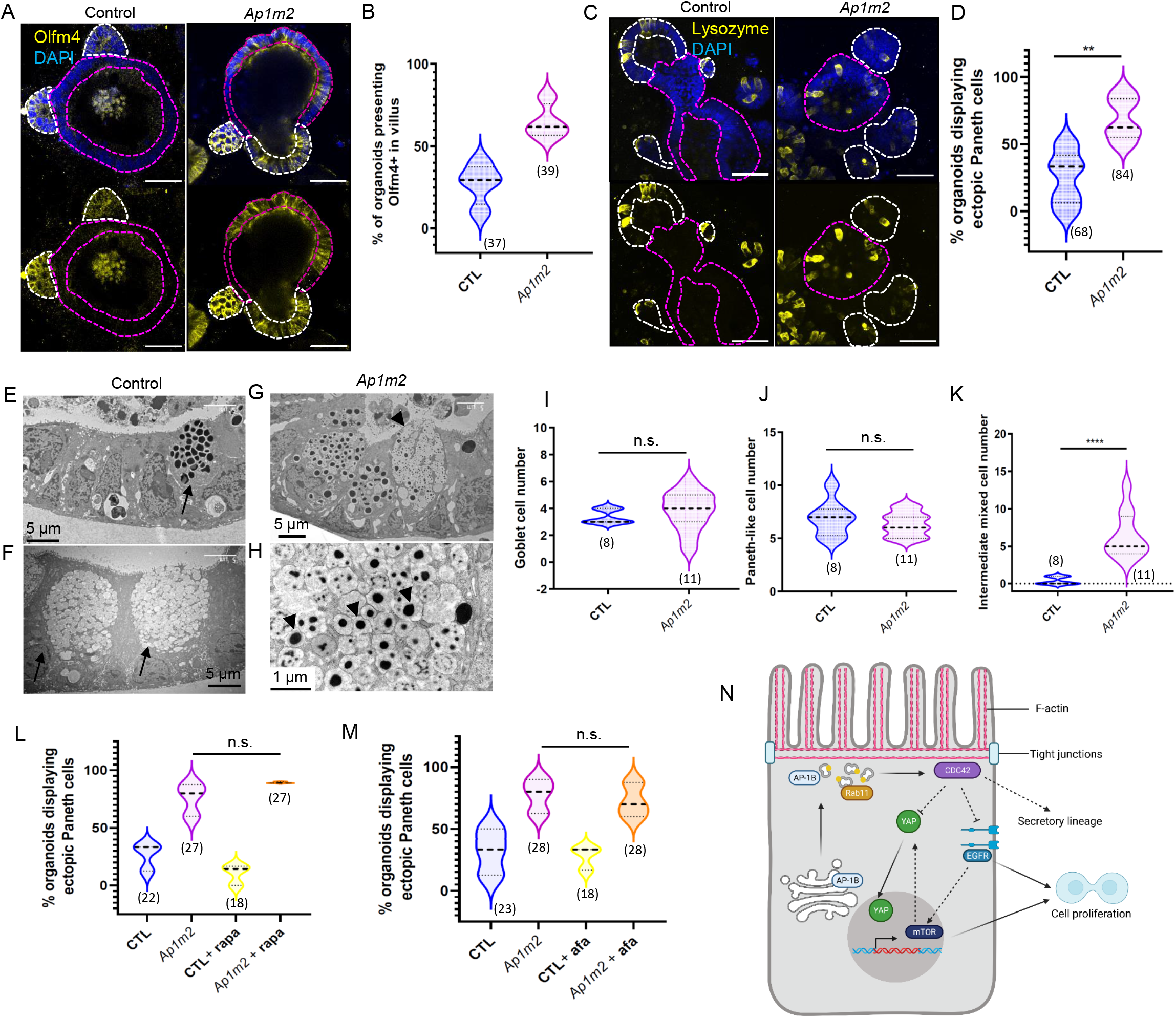
*Ap1m2* mutations affect progenitors’ distribution and secretory lineage differentiation. (A) Immunofluorescence staining of Olfm4 in Control (CTL) and *Ap1m2* organoids. (A) shows representative images and (B) the quantification of the % of organoids displaying Olmf4^+^ cells outside the crypts (N = 4 independent experiments). (C-D) Immunofluorescence staining of Lysozyme in control (CTL) and *Ap1m2* organoids. (C) shows representative images and (D) the quantification of the % of organoids displaying Lysozyme^+^ cells (N = 9 independent experiments). In (A, C) crypts and villus-like regions are delineated in white and magenta, respectively. (E-H) Secretory cells were observed by TEM, showing Paneth (E, arrow), Goblet (E, arrows) as well as intermediate mixed cells (G, H). Arrowheads show the mixed zymogen/mucigen content of intermediate cells. (I-K) Quantification of the number of Goblet (I), Paneth (J) et intermediate mixed cells (K) per organoid from 3 control (CTL) and 4 *Ap1m2* organoids, from TEM images. (L-M) Quantification of organoids displaying ectopic Paneth cells in control (CTL) or *Ap1m2* organoids treated with 200nM Afatinib (afa) (N=3 independent experiments) or1μM Rapamycin (rapa) for 3 days (N=3 independent experiments). The total number of organoids analysed is indicated in brackets, ^*^p<0,05, ^**^p<0,01; ^***^p<0,001; ^****^p<0,0001, unpaired Mann-Whitney test. (N) Model. We propose that AP-1 controls the apical organisation of Rab11 endosomes that allow the apical targeting of Cdc42, as well as inhibits the YAP and EGFR/mTOR proliferation pathways. Independently, AP-1 positively regulates the differentiation of intestinal cells towards the secretory lineage. Scale bar, 50μm.

## Material and Methods

### Materials

Afatinib and Rapamycin were purchased from Chemietek (CT-BW2992) and APExBIO (A8167), respectively.

### Organoid culture

Intestinal crypts were cultured in drops of Cultrex Reduced Growth Factor Basement Membrane Extract type 2 (BME2, 3533-010-02, Bio-techne) seeded on 12-well Greiner Cell STAR multiwell low retention plates with ENR medium (Advanced DMEM/F12 (Gibco), 10 mM Hepes (15630056), 5 ml Glutamax (35050-038), 5 ml Penicillin/Streptomycin, 1X B27 supplement (17504-044) (all from Life technologies), 1.25 mM N-acetylcysteine (A9165, Sigma-Aldrich), 50 ng/mL hEGF (AF-100-15, Peprotech), 5% R-Spondin1 and 10% Noggin conditioned media (produced as described (Sato et al., 2011)). Medium was changed every 2-3 days and organoids were passaged by mechanical disruption every 7 days. Gene mutation was induced 4 days after organoid passage during 6 days with 400 ng/ml doxycycline (D9891, Sigma-Aldrich). When indicated, full differentiation of the organoids was achieved by a concomitant treatment for 6 days with 1 mM valproic acid (P4543, Sigma-Aldrich), with the addition of 2.5 mg/mL inhibitor of WNT production-2 (IWP-2, I0536, Sigma-Aldrich) for the last 3 days, as described (Mosa et al., 2018). Absence of mycoplasma was checked regularly.

### Conditional gene knock out by CRISPR-Cas9

Organoids were transduced in a two-step manner with Edit-R lentiviral inducible Cas9 nuclease and pre-designed sgRNAs lentiviral particles (Horizon discovery, Cambridge, UK) as described earlier (Jad et al., 2022). The target sequences of the sgRNAs targeting *Ap1m2* were: 5’ACACGATGACGAAGTTGTCC3’ for sgRNA1 and 5’CTGATTAGCCGAAACTACAA3’ for the sgRNA2. Non-targeting sgRNA#1 (VSGC10215) was used to generate control organoids. Transduced organoids were selected with 5μg/ml Blasticidin (CAS9) or 3μg/ml Puromycin (sgRNAs). The % of indels was calculated using the ICE software (Conant et al., 2022) after Sanger sequencing of the organoids’ genomic DNA.

### Transmission Electron Microscopy

Organoids were collected from BME2 and fixed in Trump’s fixative at 4°C. Enhanced chemical fixation was performed in 4% paraformaldehyde/2.5% glutaraldehyde in cacodylate buffer (0.1 M, pH 7.4) overnight at 4°C. Then, organoids were incubated successively in 1% OsO4 for 1.5h and 2% uranyl acetate for 1.5h, at RT. Organoids were then dehydrated through graded ethanol solutions, cleared in acetone, and flat-embedded in Epon-Araldite mix (EMS hard formula) using adhesive frames (11560294 GENE-FRAME 65 μL, Thermo Fisher Scientific). Ultrathin 70 nm sections were cut on an ultramicrotome (UC7; Leica Microsystems) and collected on formvar-coated slot grids (FCF2010-CU, EMS). TEM grids were observed using a JEM-1400 TEM (JEOL) operated at 120 kV, equipped with a Gatan Orius SC1000 camera piloted by the Digital Micrograph 3.5 program (Gatan).

### Immunohistochemistry

Organoids were collected from BME2, fixed in 2% paraformaldehyde overnight at 4°C and washed in PBS 1X. Then, pelleted organoids were resuspended in 2 drops of Shandon Cytoblock Cell Block reagent (ThermoScientific) and embedded in paraffin wax with Excelsior ES50 during 3h. Paraffin-embedded organoid blocks were cut at 4 μm thickness, mounted on positively charged glass slides and dried at 58°C for 60 min. The slides were then deparaffinized with xylene (3×10 min) and successive 100%-75%-50% ethanol baths (5 min each) before rehydration in water for 5 min. Immunohistochemical staining was performed on a Discovery Automated IHC Stainer. First, antigen retrieval was performed using Tris-based buffer solution CC1 at 95–100°C for 48 min. Then, endogen peroxidase was blocked with Inhibitor-D/3% H2O2 for 8 min at 37°C. After rinsing, slides were incubated at 37°C for 60 min with anti-Cdc42 (1:50, ab64533, Abcam) and anti-phospho-ezrin (1:2000, ab47293, Abcam) primary antibodies. After rinsing, signal enhancement was performed using the Ventana DABMap (for horse anti-rabbit HRP) or OMNIMap (for goat anti-mouse/rabbit HRP or donkey anti-Goat HRP) detection kits (Ventana Medical Systems). Slides were then counterstained with haematoxylin (16 min) and bluing reagent (4 min) and rinsed before being manually dehydrated and coverslipped. Imaging was performed using an Hamatsu scanner (Nanozoomer 2.0RS/ C10730-12) and the images were analysed with NDP viewer.

### Immunostaining

For immunostaining on whole organoids, they were collected from BME2 and fixed in 2% PFA/PBS overnight at 4°C. For immunostaining on organoid slices, 4 μm paraffin bloc sections were mounted on slides before deparaffination in Xylene (3×10 min) and successive 100%-75%-50% Ethanol baths (5 min each). After rehydration in water for 5 min, antigen retrieval was performed with Tris-EDTA, pH8 (98°C, 40 min). Then, both organoids 4 μm slices and whole organoids in microtubes were permeabilized in PBS1X, 0,1% Tween20, 0,2% Triton (20 min at room temperature under rotation) and incubated overnight at 4°C in blocking solution (PBS1X, 0.1% tween 20, 2% FBS) with primary antibodies. Whole organoids were incubated with anti-Ki67 (1:100, 14-5698-82, Invitrogen), anti-EphB2 (1:100, AF467-SP, Biotechne), anti-YAP (1:100, 14074, Cell signalling), anti-Olfm4 (1:400, 39141, Cell signalling), anti-lysozyme (1:100, A0099, Agilent). Organoid slices were incubated with anti-phospho-ezrin (1:200, ab47293, Abcam), anti-Rab11A (1:100, 71-5300, Invitrogen), anti-Cdc42 (1:50, ab64533, Abcam), and anti-E-cadherin (1:200, 24E10, Cell signalling). Then, organoids were incubated with adequate fluorescent secondary antibodies (Jackson ImmunoResearch) during 1h at room temperature before mounting in ProLong Gold Antifade reagent (P36941, Life Technologies). F-Actin was stained using Actistain (1:150, Cytoskeleton) for 1h at room temperature. Imaging was performed on a Zeiss LSM880-Airyscan (Oberkochen, Germany) equipped with a 63X, 1.4 NA objective (Zen Black software) and images were analysed using Fiji.

### Quantification and statistical analysis

The nuclear accumulation of Ki67 and YAP was quantified by measuring the signal on a nuclear overlay obtained after nuclear segmentation using the DAPI channel in Fiji and was divided by the signal of the same overlay in the Dapi channel to correct for signal differences due to the varying distance between the cells and the objective. The apical/cytoplasmic ratio of markers was calculated by measuring the signal intensity of a line covering the apical membrane and measuring the signal intensity of an identical ROI in the underlying cytoplasm. Gaussian distribution and homoscedasticity of data were verified by the Shapiro-Wik test and the F-test, respectively. Mann-Whitney non-parametric test was performed using GraphPad software.

## Abbreviations

AP-1B: Adaptor protein-subunit μ2/μ1B
EGF: Epidermal Growth Factor
ISC: intestinal stem cells
mTOR: mechanistic target of rapamycin
YAP: yes-associated protein 1

## Acknowledgments

We thank Hans Clevers for the Noggin and R-Spondin1 expressing cells and Justine Viet for her help with CRISPR-Cas9-mediated mutations analysis. IHC and imaging were performed at the Histo Pathology High Precision (H2P2) and the Microscopy Rennes Imaging Centre (MRiC) facilities of the UMS Biosit, member of the national infrastructure France-BioImaging supported by the French National Research Agency (ANR-10-INBS-04).

## Figure legends

**Supplementary Figure 1.**
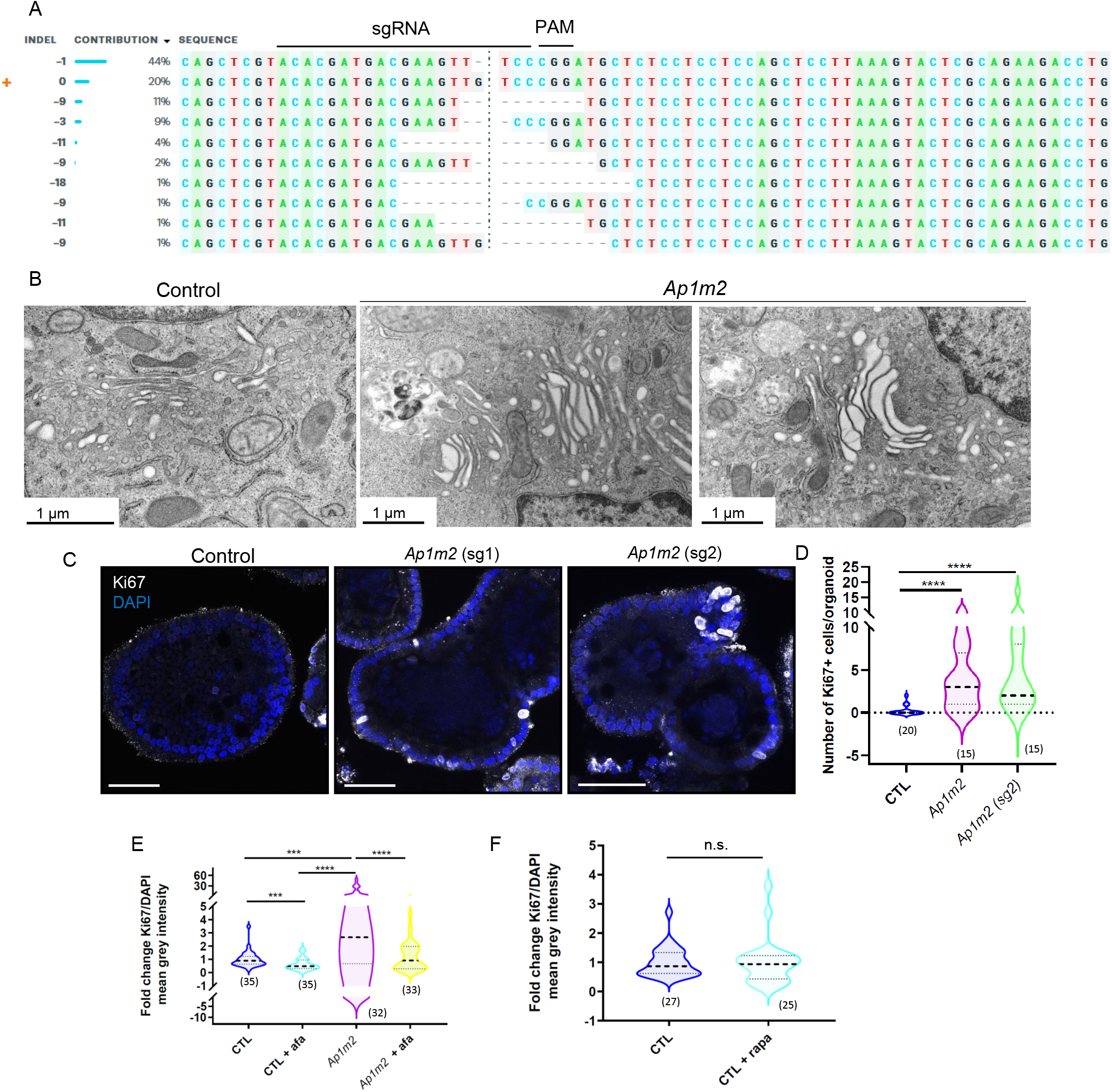
(A) ICE analysis of a genomic DNA sequencing of CRISPR-edited organoids. The sgRNA sequence (sg1) eting *Ap1m2* gene and the PAM site are indicated. (B) TEM analysis of the Golgi apparatus in control and *Ap1m2* organoids. (C-D) munofluorescence staining of Ki67 in fully-differentiated control (CTL), *Ap1m2* (sg1) and *Ap1m2* (sg2) organoids. Scale bars, 50 μm. (D) shows the quantification of Ki67+ cells/organoid. (E-F) Quantification of Ki67/DAPI ratio in organoids treated with 200 nM tinib (afa) during (N=3 independent experiments) or 1μM Rapamycin (rapa) during (N=3 independent experiments). The total number of organoids analysed is indicated in brackets. ^*^p<0,05, ^**^p<0,01 ; ^***^p<0,001; ^****^p<0,0001, unpaired Mann-Whitney test.

**Supplementary Table 1.**
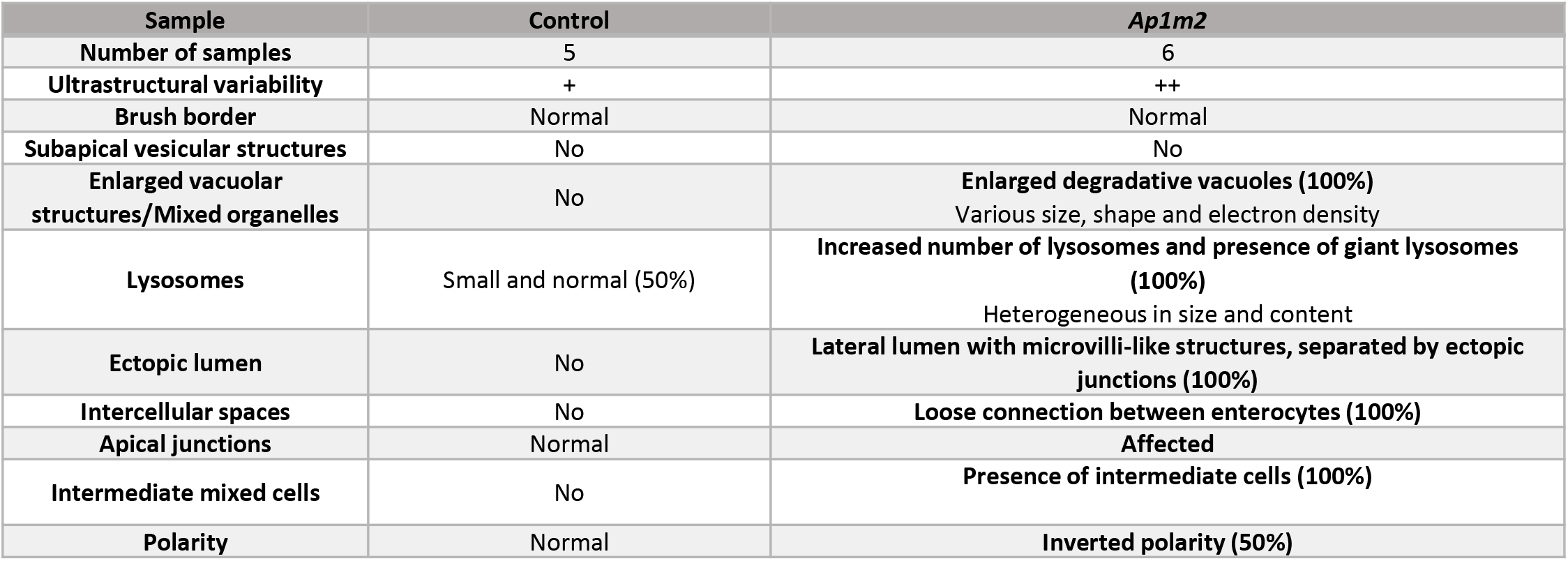
Phenotypic analysis of Control and *Ap1m2* fully-differentiated organoids by TEM. The percentages correspond to the percentage of organoids displaying the indicated phenotype.

